# Epidemiological consequences of enduring strain-specific immunity requiring repeated episodes of infection

**DOI:** 10.1101/674135

**Authors:** Rebecca H. Chisholm, Nikki Sonenberg, Jake A. Lacey, Malcolm I. McDonald, Manisha Pandey, Mark R. Davies, Steven Y. C. Tong, Jodie McVernon, Nicholas Geard

## Abstract

Group A *Streptococcus* (GAS) skin infections are caused by a diverse array of strain types and are highly prevalent in Indigenous and other disadvantaged populations. The role of strain-specific immunity in preventing GAS infections is poorly understood, representing a critical knowledge gap in vaccine development. A recent GAS murine challenge study showed evidence that sterilising strain-specific and enduring immunity required two skin infections by the same GAS strain within three weeks. This mechanism of developing enduring immunity may be a significant impediment to the accumulation of immunity in populations.

We used a mathematical model of GAS transmission to investigate the epidemiological consequences of enduring strain-specific immunity developing only after two infections with the same strain within a specified interval. Accounting for uncertainty when correlating murine timeframes to humans, we varied this maximum inter-infection interval from 3 to 420 weeks to assess its impact on prevalence and strain diversity. Model outputs were compared with longitudinal GAS surveillance observations from northern Australia, a region with endemic infection. We also assessed the likely impact of a targeted strain-specific multivalent vaccine in this context.

Our model produced patterns of transmission consistent with observations when the maximum inter-infection interval for developing enduring immunity was 19 weeks. Our vaccine analysis suggests that the leading multivalent GAS vaccine may have limited impact on the prevalence of GAS in populations in northern Australia if strain-specific immunity requires repeated episodes of infection.

Our results suggest that observed GAS epidemiology from disease endemic settings is consistent with enduring strain-specific immunity being dependent on repeated infections with the same strain, and provide additional motivation for relevant human studies to confirm the human immune response to GAS skin infection.

**Author summary:** Group A *Streptococcus* (GAS) is a ubiquitous bacterial pathogen that exists in many distinct strains, and is a major cause of death and disability globally. Vaccines against GAS are under development, but their effective use will require better understanding of how immunity develops following infection. Evidence from an animal model of skin infection suggests that the generation of enduring strain-specific immunity requires two infections by the same strain within a short time frame. It is not clear if this mechanism of immune development operates in humans, nor how it would contribute to the persistence of GAS in populations and affect vaccine impact. We used a mathematical model of GAS transmission, calibrated to data collected in an Indigenous Australian community, to assess whether this mechanism of immune development is consistent with epidemiological observations, and to explore its implications for the impact of a vaccine. We found that it is plausible that repeat infections are required for the development of immunity in humans, and illustrate the difficulties associated with achieving sustained reductions in disease prevalence with a vaccine.

## Introduction

The development of immunological memory following infection or vaccination against a particular pathogen enables a more rapid and enhanced immune response during subsequent infections. The characteristics of this immunological memory at an individual host level – such as the degree or duration of immune protection against subsequent pathogen encounters – impact epidemiological dynamics at the host population level [1,2].

Routine vaccination programs targeting pathogens comprised of a single serotype (*i.e.*, one immunologically-equivalent strain), such as the mumps and measles viruses, inhibit sustained transmission because they result in the accumulation of hosts with enduring immunological memory (herd immunity) effective against all pathogen genotypes [3,4]. For pathogens with multiple serotypes (*i.e.*, multi-strain pathogens), such as *Neisseria meningitidis* [5], poliovirus [6], *Streptococcus pneumoniae* [7] and dengue virus [8], infection by one strain may lead to an immune response that is strain-specific, providing less, if any, protection against other strains (cross-strain immunity). As a result, the link between an individual’s immune response and the accumulation of herd immunity at the host population-level can be more complex for multi-strain pathogens, posing challenges for understanding their transmission and for control [1,2,9–15].

An important human pathogen with very high strain diversity is group A *Streptococcus* (GAS), which, globally, is comprised of over 230 molecular sequence types [16] and over 290 distinct genotypes [17]. GAS generally causes infections of the skin or throat that are mild and easily treated. However, mild GAS infection can also lead to more serious invasive and immune-mediated disease with high mortality rates [18]. Hence, populations with high rates of mild GAS infections tend to also suffer from high rates of invasive disease and immune sequelae, such as acute rheumatic fever and acute post-streptococcal glomerulonephritis [18]. These GAS “hyper-endemic populations” also tend to have much higher strain diversity compared to populations with a low prevalence of GAS [19]. For example, dozens of strains of GAS are reported to co-circulate in the Indigenous communities of tropical northern Australia, where the prevalence of GAS skin infections can be as high as 45% in children, and the incidence of acute rheumatic fever is among the highest reported in the world [18,20–24].

Despite the high global burden of GAS disease [18], currently there is no licensed GAS vaccine, although there are a number in the vaccine pipeline [25]. A critical knowledge gap in GAS vaccine development is our limited understanding of how strain-specific immunity might prevent GAS infection (particularly skin infection) and, in turn, shape patterns of transmission across different populations. Epidemiological studies indicate that GAS skin infection is much less frequent in adults than in children [20,24,26,27], suggesting that people may be able to acquire enduring immunity to particular GAS strains following skin infection. However, if enduring strain-specific immunity to GAS is possible, the high rates of repeat skin infections observed in children in hyper-endemic regions [27–29] suggest that it is slow to develop. Moreover, an association between the age-related immunity to GAS and the acquisition of GAS specific antibodies suggest the need for repeated GAS exposures for enduring immunity [30]. A recent study in mice showed evidence that sterilising strain-specific and enduring immunity required two skin infections by the same GAS strain within three weeks [31]. A single infection, or two infections by the same strain that occurred greater than three weeks apart did not result in the generation of memory B cells, but rather only short-lived strain-specific immunity. An analogous mechanism of acquiring enduring strain-specific immunity from GAS skin infection in humans may be a significant impediment to the accumulation of herd immunity, particularly in populations with high numbers of circulating strains.

In this work we use mathematical modelling to determine the population-level consequences of enduring strain-specific immunity that is contingent on hosts experiencing two repeated episodes of GAS infection by the same strain [31]. We explore the effects of this immunity mechanism in the context of GAS transmission in small connected communities that are typical of Australian Indigenous populations in northern Australia, as well as other GAS hyper-endemic regions such Fiji and Samoa []. A key element of the model is that hosts can only acquire enduring immunity protecting against reinfection by a particular strain if they experience two repeated episodes of infection by this strain within a specified time interval. To the best of our knowledge, this is the first time a transmission model of any pathogen has accounted for this type of strain-specific immunity. We use the model to assess whether epidemiological observations of GAS in hyper-endemic populations are consistent with this type of immune response. We also assess the impact of one of the leading multivalent strain specific GAS vaccines on interrupting transmission in this context. Understanding generated may be crucial for predicting and understanding future population effects of GAS vaccines currently in development [25].

## Methods

In this section we describe our agent-based model of GAS transmission, the selection of model parameters based on available epidemiological studies, and our *in silico* experiments.

### Model of GAS transmission

Our agent-based model simulates the transmission of *n*(*t*) strains of GAS in a well-mixed host population (where agents correspond to hosts) of constant size *N*, in discrete time *t*. We assume the population is situated in a geographical region where *n*_max_ strains of GAS are in circulation so that 0 ≤ *n*(*t*) ≤ *n*_max_. Each strain is assumed to have on average identical transmissibility, cause infections with identical baseline average duration, and be equally distant to each other in ‘antigenic strain space’ [32] so that each strain prompts a distinct immune response in hosts.

The model tracks the age, infection and immunity status of each host through time. Changes in host infection and immunity status occur due to the clearance of infections, transmission events, and waning immunity (detailed below), and are updated synchronously at the end of each day. New susceptible individuals aged zero are introduced into the population at a per capita rate *d* to replace individuals that are lost due to natural death. We also model migration at a per capita rate of *α* (detailed below).

#### Infection

In high incidence settings, multiple strains of GAS have been concurrently detected in the same and different skin lesions of individuals [33]. Therefore, in our model, hosts can be co-infected by multiple strains. We assume that a host can have a maximum of *κ* infections at any one time (including multiple infections of the same strain), and that the susceptibility of hosts to infection decreases as the total number of infections in each host increases. These assumptions incorporate the effects of pathogen populations directly competing for space and resources within the host, or indirectly interacting via the host immune response. We calculate the relative susceptibility *r* of host *i* to an uninfected host as

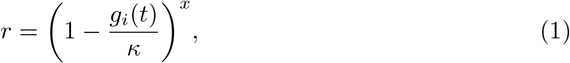

where *g*_*i*_(*t*) is the total number of infections of host *i* at time *t* and *x* > 0 is a number scaling the level of resistance to acquisition of new infections due to the competitive advantage of already established infections. Clearly, if host *i* is uninfected then *r* = 1, and if the host is at infection carrying capacity *κ* then *r* = 0.

Each day, each infecting strain will clear with probability Γ = 1 *−* exp(*−γ/s*), where 1/*γ* is the mean duration of infection of a host without prior immunity, and *s* is the expected relative duration of infection of a host compared to a host without prior immunity (detailed below). If a host has multiple infections of the same strain and this strain clears during a time step, then we assume that all infections of that strain in the host clear simultaneously.

#### Transmission

In the model, each host has on average *c* contacts with other hosts per day. The contacts of infected hosts are chosen uniformly at random from the population, and the outcomes of these contact events are then determined (*i.e.*, whether or not a transmission event occurs). We specify that transmission may only occur one-way from the infected host to their contacts. The probability of a contact resulting in transmission is *B* = *βr*, where *β* is the baseline probability of transmission, and *r* is the relative susceptibility of a host to an uninfected host (detailed above). If the infected host has more than one infection, only one of these co-infections can possibly transmit during a single contact event. For co-infected hosts, we choose one infection uniformly at random to attempt transmission. If this attempt fails, then the contact event does not result in transmission. These rules correspond to the assumption that co-infected hosts are not necessarily more infectious than hosts with a single infection. We also specify that a host may only contract a maximum of one infection per day.

With these assumptions, we can calculate the *basic reproduction number* 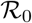, which is the expected number of secondary infections caused by a single infected host introduced into a completely susceptible host population. A pathogen is expected to cause an outbreak or become endemic in a host population if 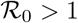. In our model, 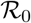 is defined as

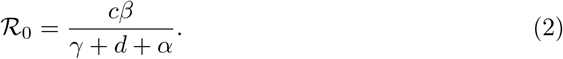

#### Immunity

Based on observations in the mouse model of GAS skin infection discussed above [31], we assume that the clearance of any host’s first infection by a particular strain confers temporary immunity. This temporary immunity has a strain-specific effect of strength *σ* (where 0 ≤ *σ* ≤ 1) and a cross-strain effect of strength *ω*_1_ (where 0 *≤ ω*_1_ ≤ *σ*) that lasts for a duration *w* for all hosts and strains.

If a host has temporary strain-specific immunity to a particular strain and is reinfected by the same strain, clearance of this subsequent infection leads to enduring strain-specific immunity that prevents reinfection by this strain and confers enduring cross-strain immunity of strength *ω*_2_ that is effective against strains that a host does not have temporary or enduring strain-specific immunity to. However, if this temporary immunity wanes, then a subsequent infection by this strain will only confer temporary immunity with the same characteristics as a first infection. This natural history of infection is summarised in Fig 1. Henceforth, we refer to the duration *w*, as the ‘maximum inter-infection interval’ that enables the development of enduring strain-specific immunity.

**Fig 1.**
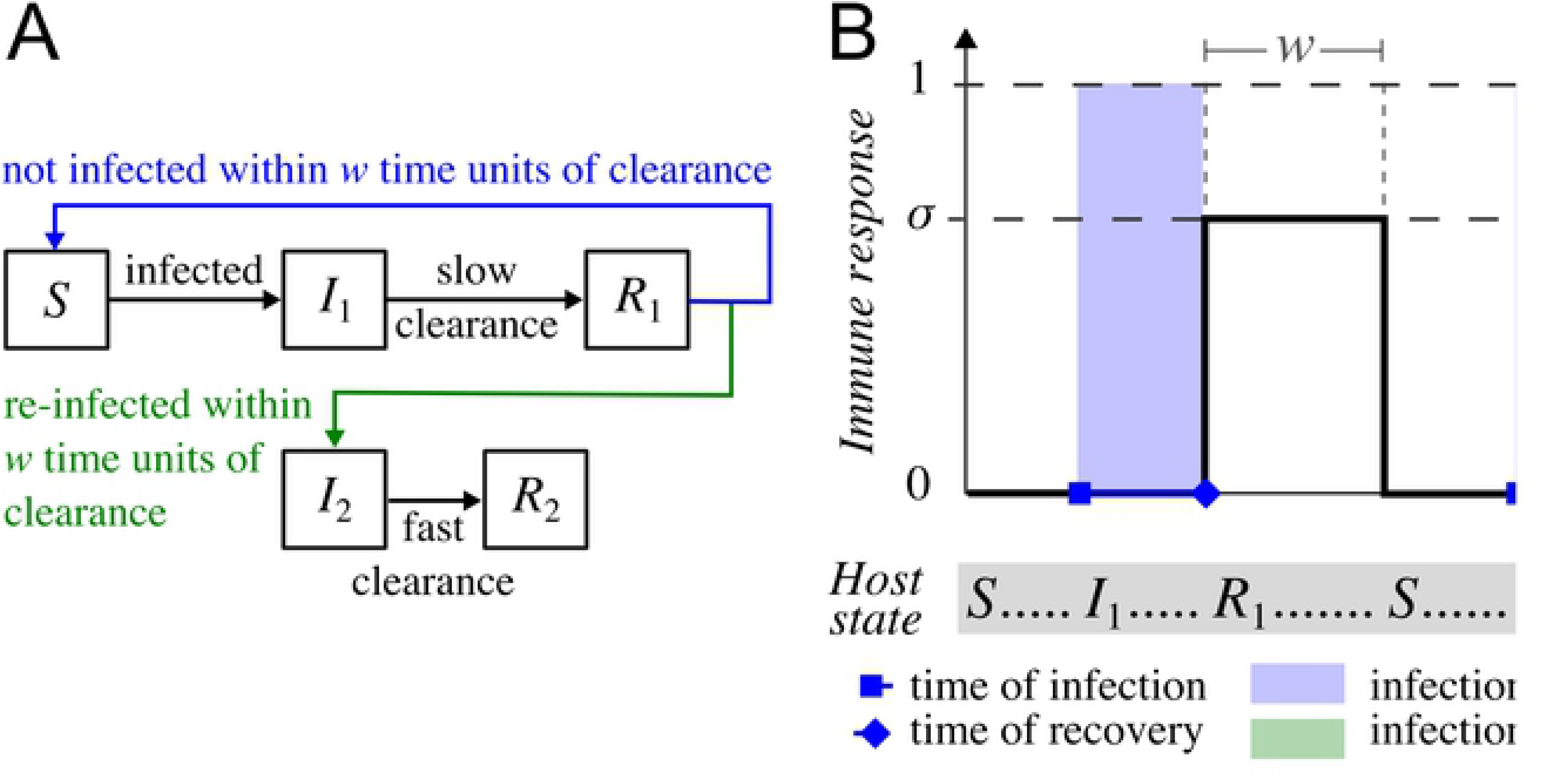
Model of the natural history of disease with respect to a single strain. A: Hosts without prior immunity to a particular strain (*S*) become infected by contacting infected hosts (*I*_1_ or *I*_2_). These infections (*I*_1_) clear at an average rate *γ* which confers temporary immunity (*R*_1_). This temporary immunity reduces the duration of a subsequent infection (*I*_2_) by a factor dependent on the strength of temporary strain-specific immunity (*σ*) if the subsequent infection occurs within a short-enough time window (the maximum inter-infection interval, *w*) from the time of clearance (green line). If infection does not occur within this time frame (blue line), then temporary immunity wanes and a subsequent infection has the characteristics of a first infection. If temporary immunity does not wane before the next infection, then the clearance of this next infection occurs faster, and confers enduring immunity protecting against further infection (*R*_2_). B: An example of a host’s immune response (solid black line) following three episodes of infection by the same strain. Here, the temporary immunity acquired following the first infection wanes before the second infection. The clearance of the second infection leads again to temporary immunity. However this becomes enduring immunity following the clearance of the third, more timely, infection. Note that the immune response is implicitly assumed to accumulate during an infection, leading to clearance (as shown by the dotted lines).

We note that in the model, it is possible for a host without any prior strain-specific immunity of a strain to experience multiple infections of a particular strain simultaneously. Due to our assumptions about strain clearance (detailed above), all infections by the same strain will clear simultaneously in the model when the host recovers from this strain, leading to a single immune response. We assume that such a clearance event only confers temporary immunity.

In the mouse model [31], the effect of the immune response was assessed by determining the number of colony forming units in skin and blood samples (bioburden) collected six days post inoculation. These showed a reduction in bioburden of approximately 90% for second infections of the same serotype compared to the first infection, provided that the second infection occurred within three weeks of the first. However, if the second infection was a different serotype, this reduction in bioburden ranged from approximately 0–30%. Our model does not explicitly represent bioburden within hosts. However, a reduction in bioburden during an infection could conceivably result in a reduced duration of infection and/or reduced infectiousness of the host. In our model, we translate the reduction in bioburden due to host immunity into a reduction in the duration of infection. Specifically, for each host *i* their expected relative duration of an infection by strain *j* compared to a host with no immunity is

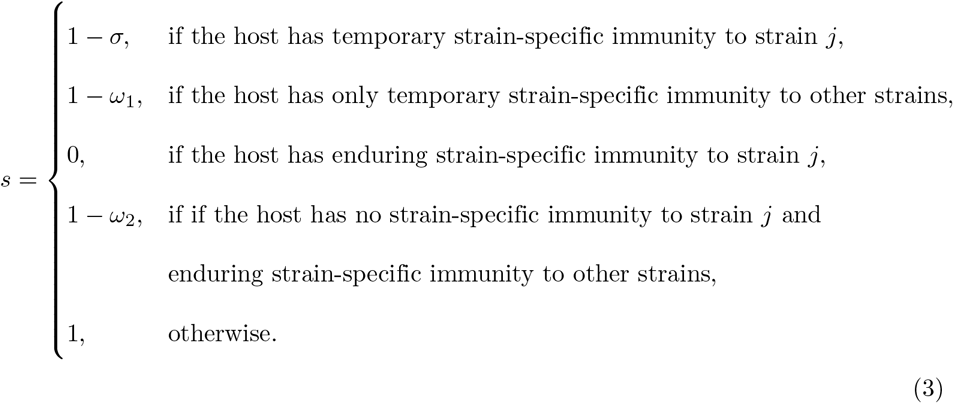

Clearly, if a host has no immunity then the expected duration of an infection is not reduced from the baseline duration 1/*γ* (since *s* = 1 in this case). If a host has temporary strain-specific immunity of a strain at the time they are infected by this strain, then the expected duration of infection is reduced according to the strength of temporary strain-specific immunity *σ* (so that *s* = 1 − *σ*). A host with enduring strain-specific immunity to a strain is essentially completely protected against infection by this strain (since a subsequent infection by this strain will have zero duration). Without strain-specific immunity to a strain, a host may still have a shorter expected duration of infection by that strain if they have either temporary or enduring immunity of other strains at the time of infection (since either *s* = 1 − *ω*_1_ or *s* = 1 − *ω*_2_ in these cases).

#### Migration

In host settings where GAS disease is hyper-endemic and where high numbers of GAS strains typically co-circulate, different strains of GAS have been observed to move sequentially through communities rather than persist indefinitely [21–24]. The introduction of novel strains and previously circulating strains into these populations is thought to be enabled by host mobility [34,35]. Therefore, in our model, each day *A* hosts (where *A* is a Poisson distributed random variable with mean *αN*) are chosen uniformly at random to be replaced by immigrants. Immigrants are assumed to have a similar immune profile to individuals in the population. This is implemented by specifying that an immigrant will have the same immune profile as an individual selected uniformly at random from the population. Immigrants may also be infected with up to one copy of infection of any strain (chosen uniformly at random from all *n*_max_ strains in the region). The prevalence of infection in immigrants is set at 10% to be consistent with the asymptomatic carriage rate of GAS across all age groups and population settings [36].

### Summary statistics

Two metrics are used to summarise transmission dynamics in our model at the population-level at time *t*: the diversity of strains *D*(*t*), and the prevalence of infected hosts *P* (*t*). We choose these summary statistics as they can be calculated from existing epidemiological data of GAS transmission [24]. Strain diversity is a measure of the total number of strains as well as how evenly strains are distributed across all infections in the host population. We calculate strain diversity using Simpson’s reciprocal index, *D*(*t*):

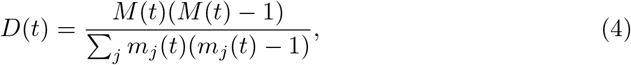

where *m*_*j*_(*t*) is the number of infections of strain *j* in the host population at time *t*, and *M* (*t*) is the total number of infections in the host population at time *t*. The prevalence of infected hosts in a host population, *P* (*t*), is calculated as

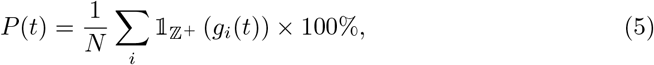

where 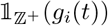 is the indicator function of the subset 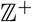 (the positive integers) of the set of all non-negative integers 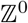 which takes the value of one when 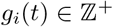 (*i.e.*, when host *i* has at least one infection) and zero otherwise. We define pathogen extinction to be the case where the prevalence *P* (*t*) = 0.

### *In silico* experimental approach

We simulate our model to (I) understand the population-level consequences of hosts requiring two episodes of infection within a given time frame to obtain enduring strain-specific immunity; (II) to determine whether epidemiological observations of GAS in an Australian Indigenous population are consistent with this type of immune response; and (III) to investigate how a targeted multivalent vaccine could potentially alter the prevalence of GAS in the Australian Indigenous context.

Since GAS is endemic in human populations, we only consider endemic transmission dynamics in our model. All simulations are run for at least 50 years to allow the epidemiological dynamics to reach a quasi-steady state where the level of immunity in the population reaches a stable level. We measure population immunity by the mean number of strains hosts in the population currently have immunity to, *Ŷ*(*t*), and we define the quasi-steady state (where *Ŷ*(*t*) is stable) as the endemic equilibrium. We also define *P*\* and *D*\* to be the endemic values of the summary statistics *P* (*t*) and *D*(*t*). These are calculated by taking the mean values of *P* (*t*) and *D*(*t*) across the previous 5 years (that is, for *t* ∈ [45, 50] years).

#### Selection of model parameters

Table 1 shows the parameters in our model and the values we considered in our simulations. Parameters were selected to reflect GAS transmission an Indigenous population of northern Australia, where GAS disease is hyper-endemic and the majority of GAS infections are skin infections [24].

**Table 1.** Parameter values

| Symbol | Description | Values / Range | Ref. |
| --- | --- | --- | --- |
| $N$ | Population size | 2500 | [24] |
| $n_{\max}$ | Number of strains circulating in region | 40 | [24] |
| $d$ | Per capita death rate (years<sup>-1</sup>) | 1/71 | [37] |
| $c$ | Mean number of daily contacts | 33 | [38] |
| $\alpha$ | Mean per capita rate of migrations (weeks<sup>-1</sup>) | 0.002 | [39] |
| $\sigma$ | Temporary strain-specific immunity strength | 0.9 | [31] |
| $\omega_1$ | Temporary cross-strain immunity strength | 0.1 | [31] |
| $\omega_2$ | Enduring cross-strain immunity strength | 0.1 | [31] |
| $1/\gamma$ | Baseline mean duration of infection (days) | 14 | [22–24] |
| $w$ | Maximum inter-infection interval (weeks) | {3,19,104,420} | [40] |
| $\mathcal{R}_0$ | Basic reproduction number | [1,10] | - |
| $\kappa$ | Maximum number of co-infections per host | 20 | - |
| $x$ | Level of resistance to co-infection | 10 | - |

The population size *N* and the number of strains circulating in the region *n*_max_ are set at 2500 and 40 respectively to be consistent with community sizes [24] and the number of strains circulating [19] among Indigenous populations of northern Australia. The mean duration of infection 1*/γ* is set at 14 days to be consistent with clinic data collected in this setting [22–24].

The number of daily contacts *c* is calculated using household contact data collected in remote Australian Indigenous communities [38]. In this setting, it is estimated that individuals make approximately 22 contacts per day on average in households. Due to a lack of data describing contact patterns outside of households in these populations, we make the assumption that an individual will have roughly half the number of contacts outside of households compared to within households (approximately 11 contacts per day), as has been assumed previously for a model of influenza transmission in this setting [38]. Therefore, we set the mean number of daily contacts *c* to be 33.

Migration patterns are not described in this settings. We set the per capita expected migration rate *α* to 0.002 per week which corresponds to an average of 5 migration events per week when the population size *N* = 2500. With the prevalence of infection in migrants set to 10%, infected migrants enter the population approximately once every two weeks, which is consistent with genomic analysis of GAS isolates collected across two Indigenous communities in Northern Australia [39].

Values for parameters relating to the effects of immunity are determined from the mouse model of GAS skin infection [31]. We set the strength of temporary strain-specific immunity *σ* to 0.9 and the strength of temporary and enduring cross-strain immunity to 0.1. These values are based on the respective observations of 90% and 0-30% reduction in bioburden in the mouse due to strain-specific and cross-strain immunity [31].

To date, 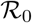 has not been calculated for GAS. We explore values of 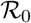 ranging from 1–10 (detailed below). For each combination of the parameters 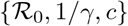 considered, the baseline transmission probability *β* is calculated using Equation (2).

#### I. What are the population-level consequences of enduring strain-specific immunity being contingent on repeat infections?

The maximum inter-infection interval *w* was estimated to be three weeks in the mouse model [31]. It is not clear how this timespan translates in humans. Based on comparisons in mice versus humans of lifespan, the time of weaning, and the age of adulthood onset, the equivalent 3-week timespan in humans could be estimated as either 104 weeks, 19 weeks or 420 weeks, respectively [40]. Therefore, to understand the population-level consequences of enduring strain-specific immunity being contingent on repeated episodes of infection of the same strain, we consider all three of these estimates for the maximum inter-infection interval *w* in humans, as well the case where *w* remains unchanged between the mouse and human, that is, with *w* = {3, 19, 104, 420} weeks.

We explore values of 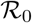 in increments of 0.5 ranging from 1–10. This range includes values of 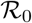 that are consistent with estimates for other pathogenic bacteria that occupy similar niches to GAS: *S. pneumoniae* [41,42] and *Staphylococcus aureus* [43] 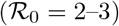. It also allows for the possibility that GAS may have a higher than expected 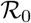 in Indigenous populations of northern Australia, where factors such as household crowding [38] and poor access to clean water [44] may increase transmissibility.

For each value of *w* and for each value of 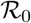 considered, we perform 80 simulations of our model. From each set of simulations, we obtain a distribution of values for the endemic prevalence *P*\* and endemic strain diversity *D*\*, from which we calculate their mean values, and 2.5%–97.5% quantiles.

#### II. Are our model outputs consistent with epidemiological data?

Next, we determine whether data simulated from our model with any of the estimates of *w* and 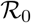 considered is consistent with epidemiological data collected in a hyper-endemic population. We compare the simulated distributions of *P*\* and *D*\* to real data collected in one Indigenous community in the Northern Territory (NT) of Australia [24]. In this study, prospective surveillance of a population of approximately 2500 people was carried out monthly over a 23 month period. Swabs were taken from the throats of all participants and any skin sores of participants and GAS isolates underwent strain typing (according to *emm* sequence). From this data we calculate the prevalence and strain diversity at each time point.

#### III. What is the potential impact of a multivalent vaccine?

A number of GAS vaccines are in the vaccine pipeline, including multivalent vaccines targeted towards serotypes associated with pharyngitis and invasive disease in Northern America and Western Europe [45]. While these targeted multivalent vaccines are predicted to provide high strain coverage in their target populations, the coverage in other populations where disease burden is much greater is predicted to be much lower [17,19]. For example, at the time of design, a leading multivalent GAS vaccine was estimated to target only 25% of the serotypes of GAS circulating in Indigenous populations of Australia, and 85-90% of serotypes in Northern America (ignoring any potential cross reactivity between serotypes) [19].

To investigate how a targeted 30-valent vaccine could potentially alter the prevalence of GAS in the Indigenous Australian context, we simulate the effects of a vaccination program consisting of routine vaccination and a one-off catch up campaign. The routine vaccination program vaccinates children when they reach one-year of age. At the commencement of the intervention, a one-off catch-up campaign vaccinates primary school-aged children in the population (aged 5–11 years). As there are no currently licensed GAS vaccines, no real-world studies to measure vaccine effectiveness exist. Therefore, we assume a best-case scenario where vaccinated hosts obtain life-long immunity from one dose of the vaccine that protects against all strains in the vaccine, and a vaccine effectiveness of 90% (which takes into account both imperfect vaccine protectiveness and imperfect program coverage).

A region-wide vaccination program will likely alter the overall prevalence of vaccine versus non-vaccine strains in the region. Therefore, strains infecting immigrants are no longer chosen uniformly at random from all *n*_max_ strains in the region. Instead, we define the probability *p*_*v*_(*t*) to be the probability that a strain infecting an immigrant will be a vaccine strain at time *t*. This is calculated as

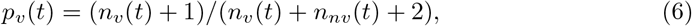

where *n*_*v*_(*t*) and *n*_*nv*_(*t*) are, respectively, the number of vaccine strains and non-vaccine strains present in the population at time *t*. This expression for *p*_*v*_(*t*) is chosen so that (1) there is a small chance that an infected migrant will be carrying a vaccine strain when there are no vaccine strains currently present in the population (since *p*_*v*_(*t*) > 0); and (2) there is a small chance that an infected migrant will be carrying a non-vaccine strain when there are no non-vaccine strains currently present in the population (since *p*_*v*_(*t*) < 1). Since we are unsure how the vaccine will affect the overall prevalence of infection, we make the conservative assumption that the prevalence of infection in immigrants remains unchanged at 10%. For every infected immigrant, if it is determined (via the probability *p*_*v*_) that their infecting strain is a vaccine strain, then this strain is chosen uniformly at random from the set of all vaccine strains. Conversely, if it is determined that their infecting strain is a non-vaccine strain, then this strain is chosen uniformly at random from the set of all non-vaccine strains. The vaccination status of any immigrants coming into the population are determined in the same way as their immune profiles – by specifying that the immune profile and vaccination status be the same as that of individuals in the population sampled uniformly at random.

We assess a range of vaccine scenarios that vary by the extent to which the 30-valent vaccine is tailored to the Australian Indigenous population context. We consider scenarios where the vaccine protects against infection by 25% of GAS strains circulating in the region (10 strains), an intermediate case where there is 50% strain coverage (20 strains), and a best-case scenario where all 30 strains targeted by the vaccine are strains that are currently circulating in the region (corresponding to 75% strain coverage). For each of these scenarios, we also explore the effect of further tailoring the vaccine to the population by choosing the vaccine strains to be the most-prevalent strains at the commencement of the intervention, as opposed to a random selection of strains (which might arise if the vaccine were tailored to another population setting).

We compare the base-line (pre-vaccine) endemic epidemiological dynamics with those calculated post-vaccine (after a further 100 years to allow the epidemiological dynamics to re-equilibrate). We also consider the short-term impact of the vaccine during the first two years of implementation. The intervention scenarios considered are further broken down into those where routine vaccination is, or is not, supplemented by the one-off catch-up campaign targeting primary school aged children.

## Results

### I. The total prevalence of infection and strain diversity are maintained by the successive reintroduction of strains

In our model, endemic transmission is characterised by continuous strain turnover rather than the persistence of individual strains over long periods of time, which is consistent with GAS epidemiological observations within endemic settings [21–24]. When individual strains appear in the population, they either fade out quickly or cause an outbreak that can last for a period of months before going locally extinct and then reappearing some time later due to a re-importation. Outbreaks of individual strains can also partially overlap, but this overlap is reduced for larger outbreaks (see panels A–B in Fig 2).

**Fig 2.**
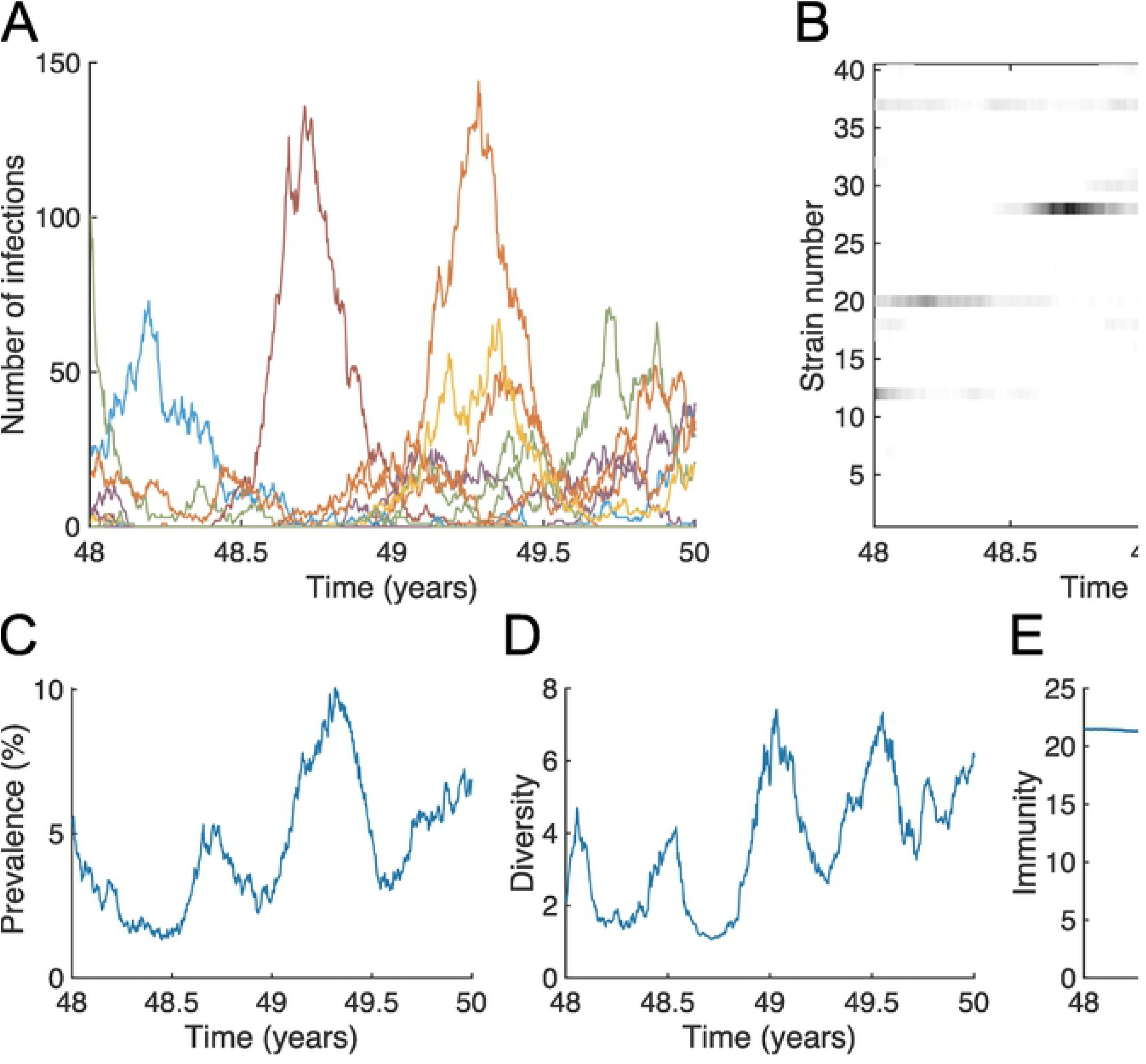
The prevalence of infection and strain diversity are maintained by the successive reintroduction of strains. Output from one realisation of the GAS transmission model over a two-year period that follows the population reaching a quasi-steady, *i.e.*, an endemic equilibrium (after 48 years). (A) The number of infections of each strain, (B) the strain distribution (with strain number on the vertical axis and shading representing the number of infections of each strain), (C) the total prevalence of infected hosts *P* (*t*), (D) strain diversity *D*(*t*) and (E) the mean number of strains *Ŷ*(*t*) that hosts have immunity to. Here, 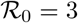, 1/*γ* = 2 weeks, *c* = 33 per day, *α* = 0.002 per capita per week, *n*_max_ = 40, *n*(0) = 35, *N* = 2500, *w* = 19 weeks, *x* = 10, *σ* = 0.9, and *ω*_1_ = *ω*_2_ = 0.1.

Despite the unstable nature of individual strains, a positive overall prevalence of infection *P* (*t*) and diversity of strains *D*(*t*) can be maintained in the population over long periods of time (see panels C–D in Fig 2) if the maximum inter-infection interval *w* and the basic reproduction number 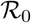 are appropriately specified (this is expanded upon below). In such cases, *P* (*t*) and *D*(*t*) oscillate around stable positive values at endemic equilibrium as individual strains sporadically appear, cause an outbreak, and then fade out. The mean number of strains that hosts are immune to, *Ŷ*(*t*), does not undergo oscillations at endemic equilibrium. Instead, it is maintained at close to a constant level (see panel E in Fig 2).

For a fixed value of the inter-infection interval *w*, increasing 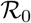 above unity initially causes both an increase in the endemic prevalence *P*\* and strain diversity *D*\* until their maxima are achieved somewhere between 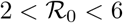 for all values of *w* considered (Fig 3). Further increases to 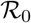 result in a slow decrease for both of these quantities. Therefore, a non-monotonic relationship exists between the basic reproduction number 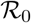 and both the endemic prevalence *P*\* and strain diversity *D*\*. For a fixed value of 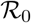, increasing *w* from the value estimated in the mouse model of infection (3 weeks), to the smallest estimate of the equivalent timespan in humans (19 weeks) has a substantial effect on reducing both the endemic prevalence *P*\* and strain diversity *D*\* for all values of 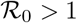 considered (Fig 3). Further increases to *w* (beyond 19 weeks to 104 and 420 weeks) correspond to increasingly smaller reductions in the endemic prevalence *P*\* and strain diversity *D*\* for all values of 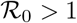 considered.

**Fig 3.**
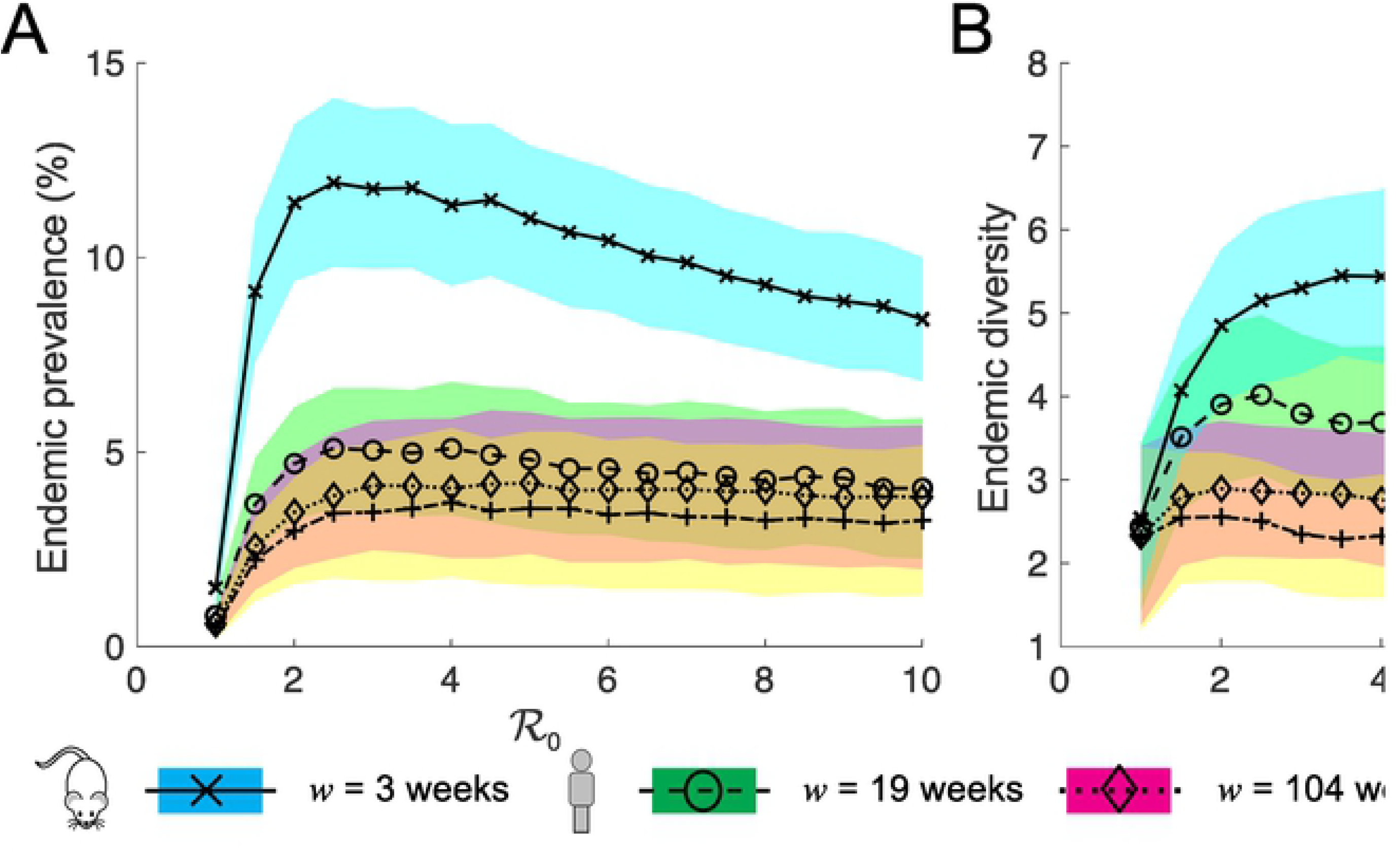
The relationship between 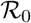 and *w*, and the endemic prevalence *P*\* and strain diversity *D*\*. The mean (lines) and the interquartile ranges (shaded regions) of (A) the total endemic prevalence of infected hosts and (B) endemic strain diversity *D*\*, from 80 simulations of the model, when the maximum inter-infection interval *w* is the value estimated in the mouse model of GAS skin infection (3 weeks), and when it is equal to three estimates of the equivalent timespan in humans (19, 104 and 420 weeks) as a function of the basic reproduction number 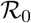 (horizontal axis). Here, 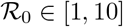, 1/*γ* = 2 weeks, *α* = 0.002 per capita per week, *c* = 33, *n*_max_ = 40, *n*(0) = 30, *N* = 2500, *x* = 10, *σ* = 0.9, and *ω*_1_ = *ω*_2_ = 0.1.

### II. Model outputs are consistent with epidemiological data collected in a hyper-endemic population

When we compare the distributions of the endemic prevalence *P*\* obtained from our simulated data and real data collected in a hyper-endemic setting we find there is only a substantial overlap of the interquartile ranges when the values of the inter-infection interval *w* is set to *w* = 19 weeks, and when the basic reproduction number 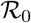 is set between 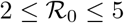 (Fig. 4A). The corresponding simulated distributions of endemic strain diversity *D*\* have substantial overlap with that of the real data for all values of *w* considered and when 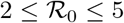. Therefore, we conclude that epidemiological data collected in a hyper-endemic population is most consistent with our simulated data generated with *w* = 19 and 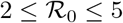.

**Fig 4.**
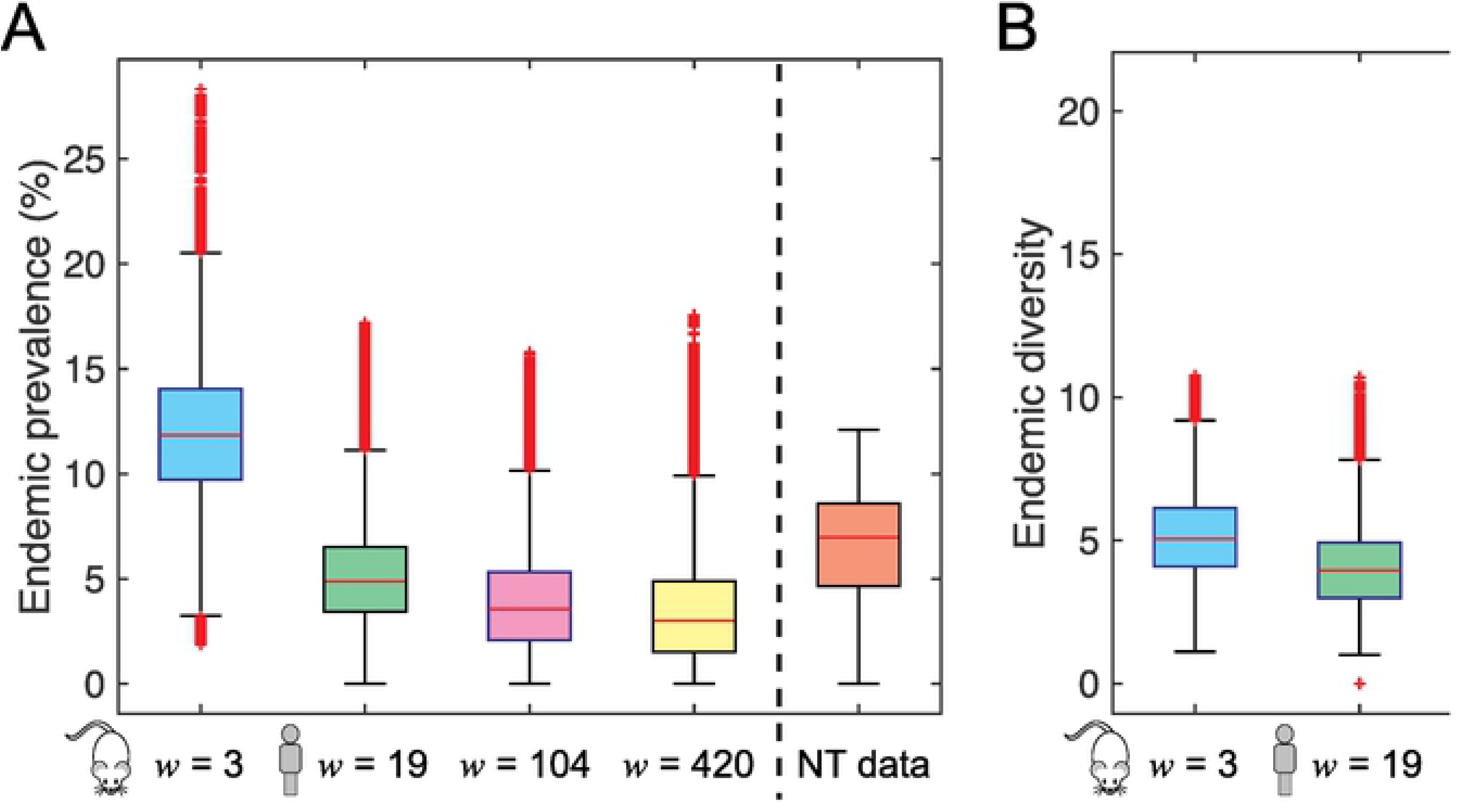
Model outputs are consistent with epidemiological data collected in a hyper-endemic population. The distribution of (A) the total endemic prevalence *P*\* of infected hosts and (B) endemic strain diversity *D*\*, from 80 simulations of the model, when the maximum inter-infection interval *w* is the value estimated in the mouse model of GAS skin infection (3 weeks), and when it is equal to three estimates of the equivalent timespan in humans (19, 104 and 420 weeks). Results are compared to population data (red) collected in one Indigenous community in the Northern Territory (NT) of Australia [24]. Here, 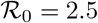, 1/*γ* = 2 weeks, *c* = 33 per day, *α* = 0.002 per capita per week, *n*_max_ = 40, *n*(0) = 35, *N* = 2500, *x* = 10, *σ* = 0.9, and *ω*_1_ = *ω*_2_ = 0.1. Similar results are obtained for 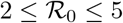 for all values of *w* considered, as evidenced by the results shown in Fig. 3, and so these results are not shown

### III. Impact of a targeted multivalent vaccine is dampened by strain replacement

When we consider the impact of a targeted multivalent (serotype-specific) vaccine on transmission, we find that the effects of the vaccine program in the short term (over the first 2 years post introduction) and in the long term (once the system reaches a new endemic equilibrium) depend on the number of distinct strains in circulation that the vaccine protects against (Fig 5). Only short-term vaccine impact is dependent on the prevalence of each vaccine strain at the commencement of the intervention, and the choice of whether or not to implement the one-off catch-up campaign.

**Fig 5.**
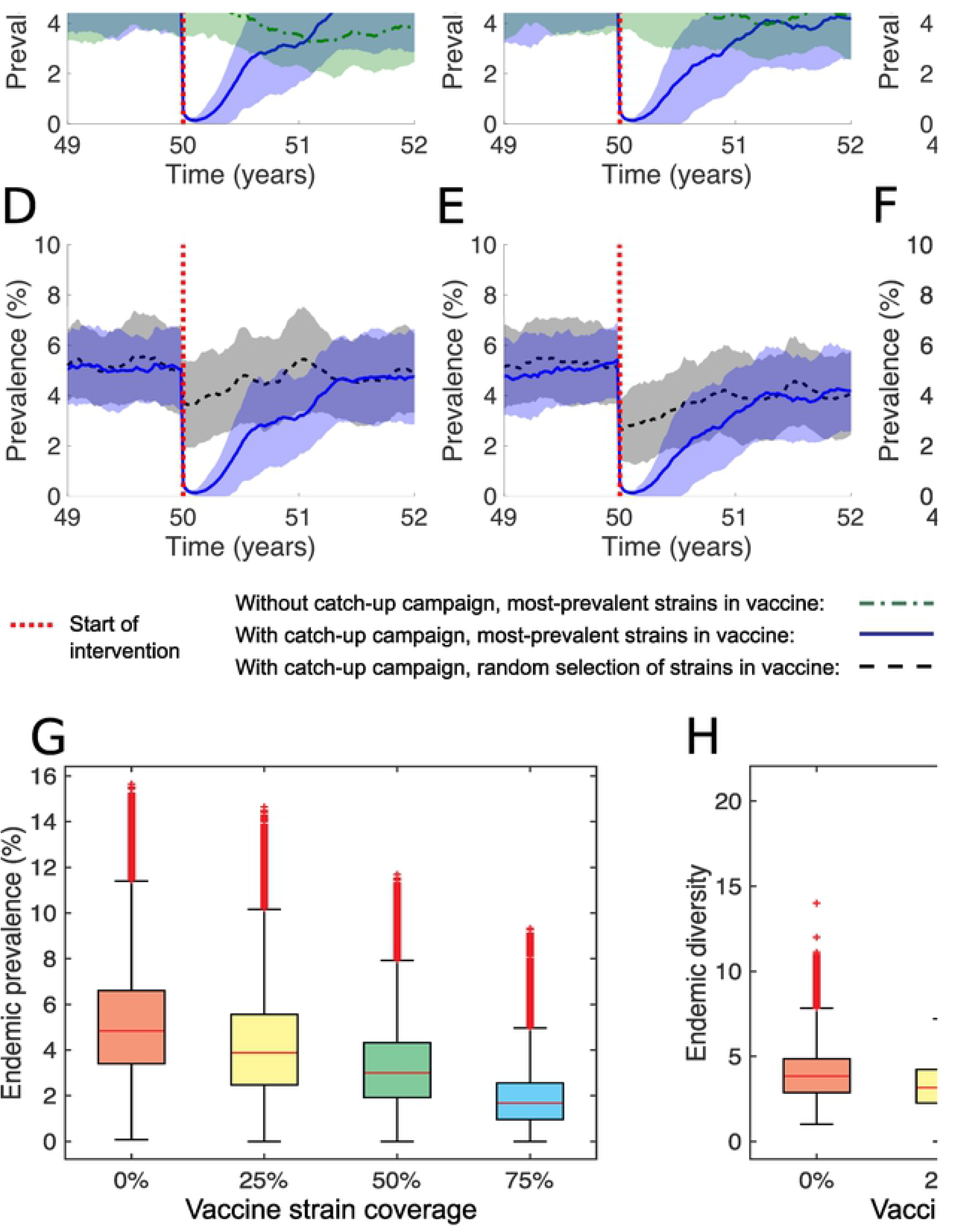
Impact of a targeted multivalent vaccine is dampened by strain replacement. (A–F) The mean (lines) and interquartile range (shaded regions) from 80 simulations of the model showing the prevalence over time, before and after the initiation of a vaccine intervention with (A,D) 25% strain coverage; (B,E) 50% strain coverage; and (C,F) 75% strain coverage. (A–C) Strains targeted by the vaccine are the most-prevalent strains at the initiation of the intervention. Scenarios with routine vaccination only (green), are compared against those where routine vaccination is supplemented with a one-off catch-up campaign (blue). (D–F) Routine vaccination is supplemented with a one-off catch-up campaign. Scenarios where strains targeted by the vaccine are the most-prevalent strains at the initiation of the intervention (blue), are compared against those where vaccine strains are randomly selected (black/grey). (G–H) The distributions of the (G) prevalence; and (H) strain diversity, calculated at endemic equilibrium pre vaccination (red boxplot) compared against those calculated post vaccination when there is (yellow) 25%, (green) 50% and (light blue) 75% strain coverage in the vaccine, when those strains targeted by the vaccine are the most-prevalent strains at the initiation of the intervention, and when there is a one-off catch up campaign as well as routine vaccination. Here, 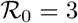, *w* = 19 weeks, 1/*γ* = 2 weeks, *c* = 33, *α* = 0.002 per capita per week, *n*_max_ = 40, *n*(0) = 30, *N* = 2500, *x* = 10, *σ* = 0.9, and *ω*_1_ = *ω*_2_ = 0.1.

Specifically, in vaccine scenarios with the one-off catch-up campaign, prevalence *P* (*t*) is quickly reduced following commencement of the vaccine program compared to equivalent scenarios without the catch-up campaign (see panels A–C in Fig. 5). This reduction in prevalence is greater when the vaccine is targeted towards the most-prevalent strains in the population at the time of the intervention, particularly for strain coverage less than 50% (see panels D–F in Fig. 5). However, prevalence rebounds in the months following the catch-up campaign to levels that are seen in equivalent scenarios without the catch-up campaign. On average, this occurs within a year when the strain coverage in the vaccine is 50% or less. When the coverage is 75%, this process takes, on average, twice as long.

In the long term, we find that the vaccine reduces the endemic prevalence *P*\* by an amount that is less than the percentage of circulating strains targeted by the vaccine. Specifically, with 25%, 50% and 75% strain coverage in the vaccine, the median endemic prevalence *P*\* is reduced by 20%, 38% and 65% respectively. The failure to fully sustain initial reductions in prevalence following vaccine introduction is due to the partial replacement of vaccine strains with non-vaccine strains, as evidenced by the corresponding small reductions in median endemic strain diversity *D*\* of 18%, 20% and 40%, respectively.

## Discussion

Incomplete understanding of the the immune response to GAS infection in individuals and the development of herd immunity in host populations represents a key barrier to the development of a globally effective GAS vaccine. Current consensus is that the immune response to GAS infection is largely strain (serotype)-specific [25]. Recent evidence in a murine model of GAS skin infection raises the possibility that the longevity of this immune response may be contingent on individuals experiencing a repeat episode of infection by the same strain within a narrow time window [31]. As yet, there is no direct evidence for an analogous immune response to GAS infection in humans.

### Indirect evidence for immunity being contingent on repeat infections

The results of our mathematical modelling study indicate that epidemiological observations of GAS infections in a population with high rates of GAS disease are consistent with enduring strain-specific immunity being contingent on repeated infection with the same strain. Both epidemiological observations [21–24] and the data simulated from our model with a sufficiently long maximum inter-infection interval *w* are reflective of there being a continuous turnover of GAS strains in the population rather than individual strains persisting over long periods of time (see Fig 2). In our model, this strain cycling is enabled by (1) infected hosts migrating into the population and triggering outbreaks of new or previously-circulating strains; (2) the accumulation of hosts with enduring immunity which causes these strains to go locally extinct; and (3) the loss of sufficient herd immunity due to the continual influx of susceptible hosts into the population (through birth and migration) which eventually allows a future reimportation to trigger another outbreak. We found that our model best matches real epidemiological data when the maximum inter-infection interval *w* is 19 weeks, and if the basic reproduction number 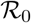 is between 2 and 5.

An alternative hypothesis of the immune response to GAS skin infection is that enduring strain-specific immunity can be acquired through the clearance of a single infection. Mathematical models of other multi-strain pathogens that incorporate this type of immune response can also exhibit high strain turnover in host populations and result in a non-monotonic relationship between 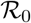 and the endemic prevalence [1,46], similar to what is observed in our model. However, this hypothesis precludes individuals experiencing repeated infections by the same GAS strain, which has been observed in children in high-incidence settings [22]. Another alternative hypothesis is that skin infection can never lead to enduring strain-specific immunity, but only temporary strain-specific immunity, thus allowing repeat infection by the same strain once immunity has waned. Future modelling work could consider whether there are conditions under which such a model is also consistent with GAS epidemiological data collected in high-incidence settings.

### Epidemiological consequences of immunity being contingent on repeat infections

Our study demonstrates the broader epidemiological consequences of enduring strain-specific immunity being contingent on repeated episodes of infection. Pathogen transmissibility has competing effects on the likelihood of hosts acquiring enduring immunity in our model, which leads to a complex relationship between transmissibility and prevalence.

Increasing the basic reproduction number 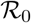 from small values initially corresponds to a rise in the endemic prevalence of infection *P*\* due to increased transmission. This increase in *P*\* continues until transmission reaches a critical level whereupon it becomes more feasible for hosts to encounter the same strain twice within the required time window *w* and acquire enduring immunity. In this regime, further increases to 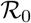 correspond to increased levels of herd immunity that eventually lead to reductions in the endemic prevalence *P*\* for further increases to 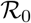. However, these further increases to 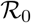 correspond to increasingly smaller reductions in the endemic prevalence *P*\*, possibly because the reduction in duration of outbreaks of individual strains (which coincide with increases to transmissibility) make it more difficult for hosts to experience multiple episodes of infection of the same strain during a single outbreak.

A possible consequence of a non-monotonic relationship existing between 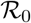 and the endemic prevalence is that interventions designed to reduce 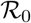 (*e.g.*, via social interventions to improve household crowding or access to healthcare or running water) may lead to different outcomes in populations characterised with different baseline 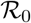. For example, an intervention that leads to a substantial reduction in prevalence in one population may lead to very little change or even an increase in prevalence in a different population that has a higher baseline 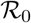.

### Short and long-term benefits of tailoring a multivalent vaccines to target populations

Our study also demonstrated how our model can be used to interpret and predict the effects of a targeted multivalent-vaccine intervention in a high-incidence setting. A key determinant of long-term vaccine impact is the number of strains that the vaccine protects against that are circulating in the greater geographic region of the population. The greatest long-term reductions in prevalence occur when all strains in the vaccine are those in circulation, indicating the importance of customising a multivalent vaccine to particular host settings, or incorporating more conserved antigens with multivalent formulations [17].

Nevertheless, the high strain turnover that characterises transmission is likely to limit the long-term effectiveness of a targeted multivalent vaccine that does not protect against *every* strain in circulation. In our model, the replacement of vaccine strains with non-vaccine strains occurred within a few years of the implementation of the vaccine intervention. This occurred even when there was 75% vaccine strain coverage, and following significant short-term reductions in prevalence. While it may be the case that initial reductions in prevalence following the introduction of the vaccine cannot be sustained, it may be possible for long-term benefits to arise if the vaccine is rolled out in combination with other interventions designed to reduce transmission. It will be crucial to conduct surveillance for a number of years following vaccine introduction to evaluate short- and long-term vaccine impact.

### Limitations and future work

In our model of GAS transmission in an Indigenous population of northern Australia, we assume that all GAS infections lead to the same type of immune response – that which was was observed in the mouse model of GAS skin infection [31] – since the majority of mild GAS infections in this setting are skin infections [23,24]. However, in lower incidence settings, current consensus is that GAS causes throat infections more frequently than skin infections [18]. Furthermore, GAS can also be carried in the nose and throat of hosts without symptoms, and, less frequently, cause invasive disease [18]. It is not clear whether these other types of GAS infections cause an analogous immune response. If so, future modelling work could consider transmission and the effect of interventions in populations where other or multiple types of immune responses to GAS infection occur.

We have assumed that all GAS strains in the model have identical epidemiological characteristics. Further empirical work is needed to determine the validity of this assumption. Given that all strains share the same ecological niche, any differences in the competitive ability of strains will likely alter the level of strain diversity that can be sustained over short and long timescales in populations [47]. Furthermore, perturbations to pathogen population structure through the implementation of a vaccine targeting a subset of strains is likely to also depend on the epidemiological characteristics of targeted strains relative to non-targeted strains [47].

There are parallels between our simulation results and observed responses to the multivalent vaccines targeting another highly diverse human pathogen, *S. pneumoniae*. *S. pneumoniae* has over 90 different serotypes, and the multivalent pneumococcal conjugate vaccines (PCVs) targeted the most prevalent *S. pneumoniae* serotypes responsible for severe disease in different populations. The response to the PCVs varied across subgroups within these populations [48,49]. However, generally there was a decrease in detection of vaccine strains and an increase in detection of non-vaccine strains following the implementation of PCV programs [50]. This is speculated to be due, in part, to strain replacement [50], similar to what occurred in our simulations. However, evolutionary factors such as serotype switching [51] and selection dynamics associated with the accessory genome, which remained relatively unchanged pre and post the implementation of the PCVs [52], may also have played a role in the observed vaccine response. Future work could consider exploring similar factors in the context of a GAS vaccine by incorporating evolutionary dynamics, such as mutation and recombination, into our model.

## Acknowledgments

We thank Jonathan Carapetis and Ross Andrews for contributions to data collection.

